# Top-Down Scoring of Spectral Fitness by Image Analysis for Protein Structure Validation

**DOI:** 10.1101/2025.05.05.652267

**Authors:** Benjamin D. Harding, Barry DeZonia, Rajat Garg, Ziling Hu, Frank Delaglio, Timothy Grant, Chad M. Rienstra

## Abstract

Nuclear magnetic resonance (NMR) spectroscopy is a powerful technique for protein structure determination, but traditional approaches require extensive manual assignment of hundreds to thousands of resonances. Here we present NMRFAM-BPHON, a novel “top-down” approach that treats experimental NMR spectra as continuous grayscale images and quantitatively scores the agreement with simulated spectra generated from candidate protein structures. This method does not require complete resonance assignments, though it can incorporate experimental chemical shifts when available to improve performance. The simulated spectra are generated from postulated resonance assignments, which can be derived either from empirical database predictions, direct interpretation, or a hybrid combination. BPHON employs a physics-based approximate polarization transfer model to predict cross-peak intensities from the internuclear distances in the decoy structure, and models the peak lineshapes using empirical, bulk T_2_ relaxation rates and literature values for scalar couplings. The resulting simulated spectra are scored relative to the experimental data by normalized cross correlation, yielding a fitness score between 0 and 1. We demonstrate BPHON’s ability to discriminate structural models, particularly in the case of ^13^C-detected magic angle spinning solid-state NMR spectra. The software is packaged with a user-friendly graphical user interface for ChimeraX, enabling advanced NMR analysis accessible without requiring extensive manual analysis.

## INTRODUCTION

Proteins carry out complex biological functions that are essential to life and a deeper understanding of protein structures is critical to strengthen our understanding of their mechanisms. NMR spectroscopy offers unparalleled insights into protein structure and dynamics at atomic resolution, providing critical information for understanding biological function of microcrystalline^1, 2^, membrane^3, 4^, and fibrous^5^ proteins. However, the power of NMR comes with significant analytical challenges. Traditional NMR structure determination relies on the “bottom-up” approach: assigning thousands of individual resonances through sequential correlations and interpreting each peak as a discrete entity. This process typically requires collecting multiple three-dimensional (3D) datasets over days to weeks, followed often by months of resource-intensive manual interpretation by NMR specialists.^6^ While this methodology is well-established and reliable for small proteins, it faces substantial limitations when applied to larger systems.

The conventional bottom-up approach has been continuously refined since its introduction four decades ago,^7^ and automated workflows are available for solution NMR of relatively small proteins.^8, 9^ In many cases manual interpretation steps are required, including processing from time to frequency domain,^10^ peak picking, identifying sequential correlations, mapping the assigned fragments to the protein sequence uniquely, and iteratively interpreting leftover resonances that do not have sufficient sensitivity to exhibit all expected correlations. This process has been automated^11–13^, but most programs of this type follow the same logic as an expert and are prone to failure in cases where peaks are missing or overlapped, as is often the case in spectra (whether solution or solid-state NMR) of large proteins. The bottom-up approach continues to be improved with better approaches to peak detection, Monte Carlo or Bayesian approaches to the statistical analysis, and improvements to semi-empirical shift prediction.^14–17^ Nevertheless, it suffers from the fundamental limitation of treating spectra as collections of individual peaks, often without explicit consideration of the peak intensities. This works well for small (<20 kDa) proteins studied by solution NMR because most cross-peaks in the 3D spectra are uniquely resolved. However, for larger proteins with longer rotational correlation times, the solution NMR signals exhibit broader linewidths and variations in intensity, complicating this procedure. Moreover, solid-state NMR (SSNMR) methods can enable data collection for much larger proteins utilizing standard labeling approaches and pulse sequences, yielding much larger numbers of cross-peaks, greater variations in peak intensities, and sometime more overlap in the 2D spectra. These considerations make the manual procedures impractical to implement in a time-efficient manner and lead to failures of standard automated assignment procedures to reproducibly converge.

“Top-down” approaches offer an alternative paradigm for NMR structure determination, starting with hypothesized protein structures and identifying the model that best matches the experimental data. CS-ROSETTA represented an early success in this direction, using assigned chemical shift data to select protein fragments from the Protein Data Bank (PDB) as well as a standard ROSETTA Monte Carlo assembly and relaxation methods.^18^ However, CS-ROSETTA requires initial resonance assignments, which remain the primary bottleneck in NMR structure determination, especially for asymmetric assemblies, membrane proteins, fibrils and large multi-domain enzymes. Our previously developed Comparative Objective Measurement of Protein Architectures by Scoring Shifts (COMPASS)^19^ methods attempted to overcome this limitation by utilizing a directed, modified Hausdorff distance geometry as the scoring function, eliminating the need for experimental assignments. While COMPASS demonstrated promise in test cases, its requirement for well-resolved spectra that can be evaluated as discrete, individual cross-peaks has limited its practical utility. When spectra exhibit such high quality, resonance assignments are often straightforward and the bottom-up approaches work well, and so COMPASS offers a relatively incremental benefit in many important use cases.

To address these persistent challenges, we have developed a fundamentally different approach that aligns with how experts analyze complex NMR spectra in challenging cases.^5, 20^ Expert interpretation frequently involves iterative assessment of partially overlapped signals and careful consideration of relative peak intensities to differentiate between various types of correlations (e.g., intra-residue v. inter-residue). This nuanced analysis, which is not explicitly incorporated into automated assignment procedures, proves essential for assigning resonances and solving structures of asymmetric assemblies^21^, fibrils^20^, and other large complexes.^22^ Capturing this information requires simulating multidimensional NMR data with quantitative calculations of the spectral intensities, homogeneous linewidths, and fine structure arising from scalar couplings (whether or not resolved visually in the spectra).

Since the experimental spectra inherently contain this rich information content—including peak positions, intensities, widths and shapes— more accurate and precise models of these features in the simulated decoy spectra should yield improved agreement with experimental spectra and depend more strongly on protein tertiary structure. Thus, the discriminating power of the scoring function exhibits a greater dynamic range and is better able to identify decoy structures that best match the raw, experimental data. For this approach to succeed, developing a quantitative model that accurately predicts spectra from candidate structural models is essential.

Here we present such a model, evaluate the validity of key approximations intended to accelerate computational efficiency, and benchmark performance with protein data sets. We utilize a combination of semi-empirical chemical shift prediction (SHIFTX2^23^), using sequential, site-specific resonance assignments when available; the two sources of hypothesized assignments can be combined. We then evaluate the performance of two first principles polarization transfer theories that predict cross-peak intensities according to internuclear distances and chemical shift ranges. We build the lineshape model with canonical scalar coupling values and bulk relaxation rates that can be measured quickly and reliably for 1D spectral series, even for very large proteins or samples in limited quantities. These tools permit decoy 2D spectra to be computed from postulated protein structures rapidly and with high accuracy. We then score the agreement with the experimental spectra, treating the spectra not as discrete data points but as images, leveraging analysis tools commonly employed in the imaging and microscopy fields such as zero normalized cross correlation (ZNCC). The result is a score between 0 and 1 for in-phase spectra and −1 and 0 for out-of-phase spectra, which objectively and quantitatively reports upon similarity between unassigned simulated and experimental NMR protein spectra.

To make this methodology broadly accessible to the scientific community, we also report a graphical user interface (GUI) that is compatible with the widely used molecular visualization software ChimeraX.^24–26^ Our software, NMRFAM-BPHON (or BPHON for short), facilitates importing models and experimental data, computed simulated spectra, scoring agreement and visualizing the results graphically. Named after the Greek myth hero Bellerophon, BPHON is designed to be user friendly, enabling structural biologists without extensive NMR training to analyze complex spectra data without requiring extensive training in the interpretation of multidimensional NMR spectra, and without the need for detailed understanding of underlying NMR theory and data processing. Additionally, BPHON provides a platform for experts to incorporate more sophisticated models for specialized applications beyond proteins, expanding the utility of NMR in structural biology and chemistry.

## RESULTS AND DISCUSSION

### BPHON Workflow

BPHON quantitatively evaluates how accurately protein structures represent experimental NMR spectra. The workflow requires three inputs: (1) protein structures; (2) a single, unassigned experimental spectrum (typically a ^13^C-^13^C 2D correlation spectrum), and (3) a resonance list (experimental, predicted or hybrid). The output is a normalized similarity score between 0 and 1 for each model, with higher score indicating better agreement. To illustrate this process, we examined the microcrystalline protein GB1 (2LGI, Figure 1A). From this structure, BPHON simulates a ^13^C-^13^C spectrum by applying a spin physics model to predict the cross-peak intensities as a function of mixing time, so that the simulated spectrum (Figure 1B) reports directly upon spatial proximity between ^13^C atoms in the protein. Estimated relaxation properties and known scalar couplings are also implemented to provide lineshapes which depend upon the structural heterogeneity of the sample. The simulated (Figure 1B) and experimental (Figure 1C) spectra, represented as grayscale images, are then scored using the image analysis algorithm zero-normalized cross correlation (ZNCC). This approach yields a quantitative similarity measure that integrates all spectral features—peak positions, intensities, lineshapes, and overall patterns—into a single comprehensive score. For the 2LGI model, this analysis produced a best BPHON score of 0.848, indicating exceptional agreement between the structural model and experimental data (Figure S1). In the subsequent sections, we assess the requirements for each step of the modeling process in order to achieve this level of agreement.

**Figure 1.**
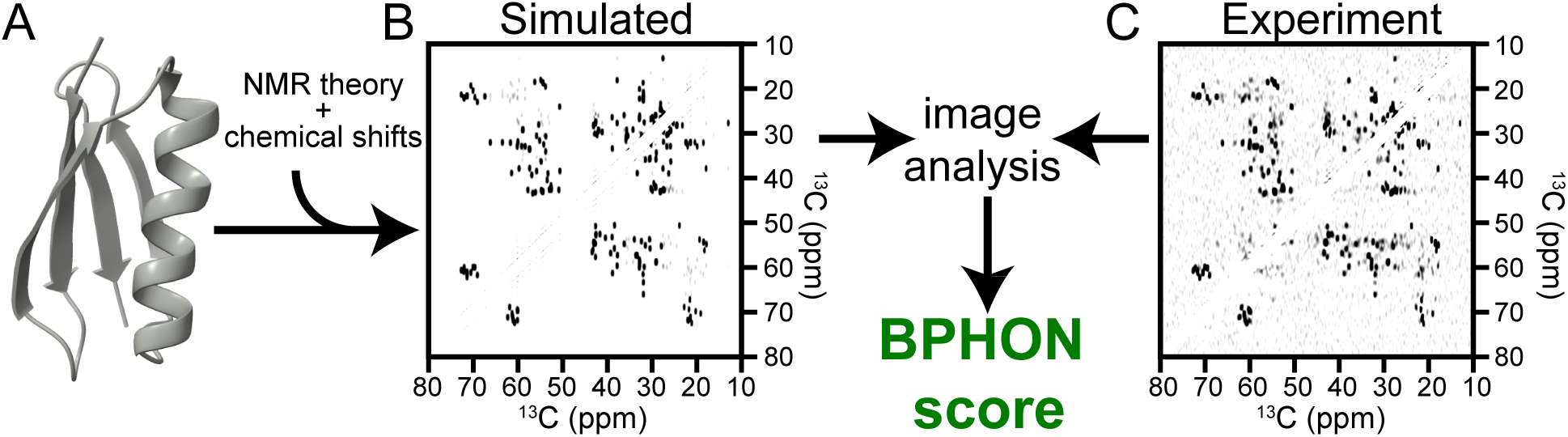
BPHON workflow. A) A protein structure of microcrystalline GB1 (2LGI) is used to simulate a ^13^C-^13^C spectrum (B) and scored against an experimental spectrum (C) using a zero-normalized cross correlation algorithm to yield a BPHON score.

### Benchmarking BPHON

The accuracy of BPHON’s simulations relies on several integrated components, including algorithms for predicting the peak positions, peak intensities, and lineshapes. Additionally, the final BPHON score is influenced by spectral processing parameters, the chemical shift accuracy, and the underlying structural model. In this section, we systematically benchmark each component and quantify their respective contributions to the overall BPHON scoring performance.

The benchmarking follows a stepwise evaluation protocol. First, we assess the accuracy of the chemical shifts used to simulate spectra: comparing predicted (SHIFTX^23^), database-derived (BMRB^27^) and experimentally determined values. Second, we contrast a simplified binary (step function) polarization transfer model with a kinetic rate equation-based approach that incorporates distance-dependent intensities. Third, we then implement amino acid-specific T_2_ relaxation rates and literature-derived scalar couplings to generate authentic lineshapes, demonstrating their substantial impact on score improvement. Fourth, we analyze how spectral processing parameters, particularly those affecting resolution and sensitivity, influence scoring outcomes. Finally, we evaluate the performance of hybrid resonance lists that combine experimental and predicted chemical shifts in various proportions and demonstrate BPHON’s ability to discriminate between alternative structural models of GB1. This comprehensive benchmarking establishes quantitative performance metrics and optimal parameters for applying BPHON to more challenging protein systems, including membrane proteins and large macromolecular assemblies.

The first step to establishing a baseline of BPHON scores is to measure the BPHON score of decoy spectra calculated with a predicted (SHIFTX2), BMRB (17810), and experimental resonance list, where the experimental resonance list is a refined set of values starting from the BMRB resonance list as initial guesses to guide assignments, but also considering the small changes that can arise due to differences in magnetic field, MAS rate and temperature. For this first stage of assessment, we use the simplest assumptions for the other aspects of the model: binary polarization transfer model where a peak intensity is either 1 or 0, and no T_2_ relaxation data or scalar couplings to calculate lineshapes. Like the COMPASS method, the principal diagonal and spinning sidebands were removed.^19^ Three ^13^C-^13^C spectra were simulated using resonance lists derived from SHIFTX2, BMRB (17810), and an experimental resonance lists assuming 50 ms of homonuclear recoupling mixing time (Table S1) and scored against an experimental ^13^C-^13^C (DARR^28^, 50 ms) spectrum collected at 14.1 T, with 26.6 kHz MAS rate. The BPHON scores for the spectra simulated with SHIFTX2, BMRB, and experimental resonance lists were 0.408, 0.489, and 0.563, respectively (Figure 2A). As expected, the experimental resonance list yields a simulated spectrum (Figure 2B) with the highest score.

**Figure 2.**
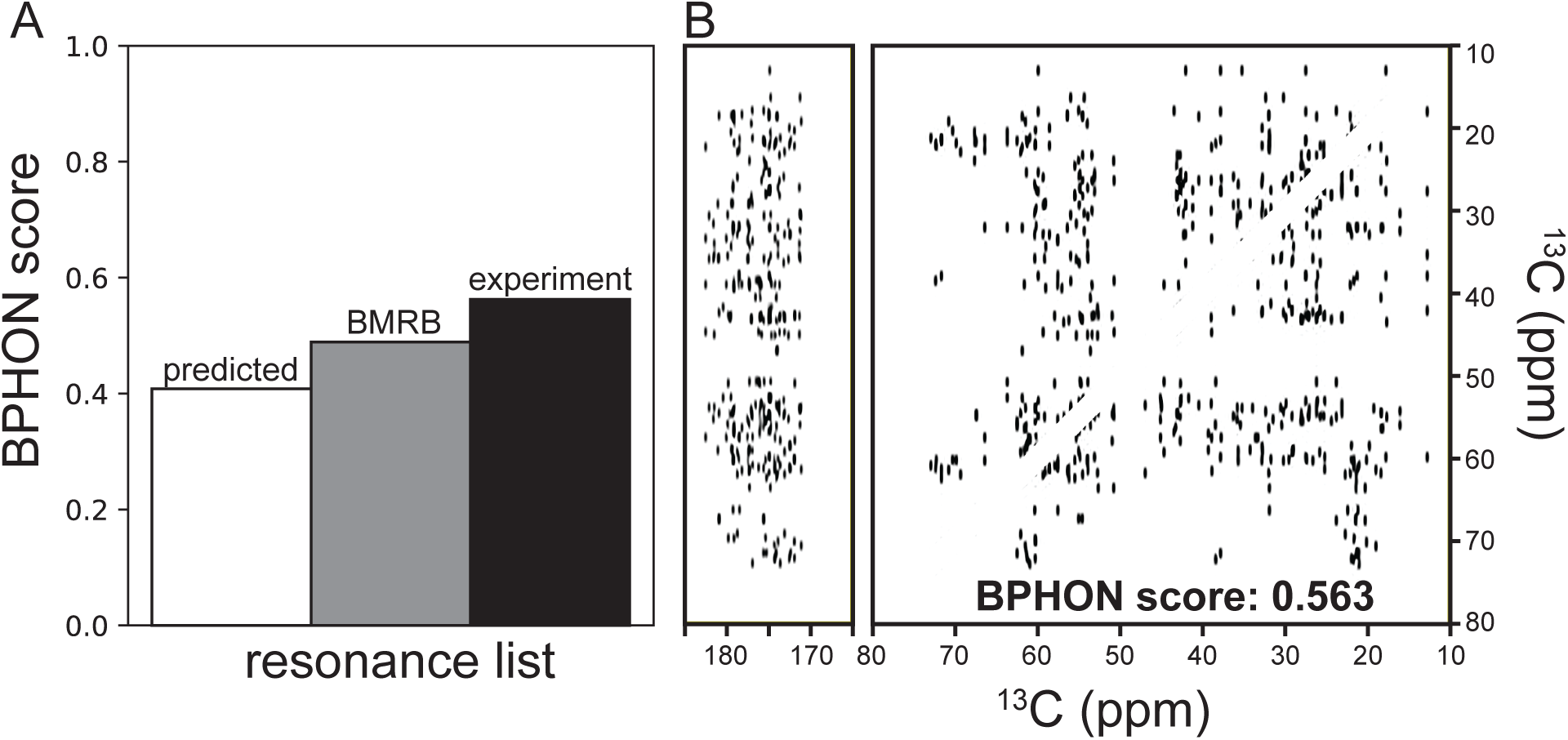
Scoring resonance list accuracy. A) histogram of BPHON scores when simulated spectra were calculated with predicted (white, SHIFTX2), BMRB (17810, gray), and experimental (black) resonance lists. B) The carbonyl and aliphatic simulated spectra of a ^13^C-^13^C spectrum simulated with the experimental chemical shifts. Simulated spectra did not employ a spin physics model to calculate peak intensities or T_2_’s and scalar couplings to provide lineshapes. Simulated spectra were scored against an experimental ^13^C-^13^C (DARR, 50 ms) spectrum collected at 14.1 T spinning at 26.6 kHz. Simulated and experimental spectra were processed with a sine bell offset of 59° in both dimensions.

Next, an improved polarization transfer model was employed to calculate peak intensities and replace the binary model. The equation used to calculate cross-peak intensity as a function of homonuclear recoupling mixing time is modified from Perras^29^:

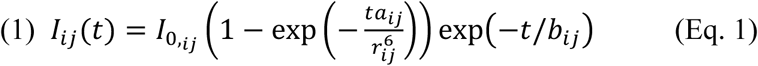

where *I_i,j_* is the intensity of the signal between spins *i* and *j*, *I_0,i,j_* is the initial magnetization scaling factor, *t* is the homonuclear recoupling mixing time (s) employed for ^13^C-^13^C spectra, *r* is the distance (Å) between two ^13^C nuclei in the protein, and *a* and *b* are independent variables used to fit experimental data to calculate peak intensity. Equation 1 and variables *I_0,i,j_, a,* and *b* were fit to nine experimental ^13^C-^13^C spectra (DARR, t_mix_ = 12.5, 25, 50, 100, 150, 200, 250, 375, 500 ms) (Figure S2). The variables that describe these parameters are presented in Table S2 and are used to calculate ^13^C-^13^C spectra using SHIFTX2, BMRB (17810), and the experimental resonance list. Consistent with the score trend in Figure 2A, the simulated spectrum calculated with the experimental resonance list has the highest score (0.737), followed by the BMRB (0.641), followed by the predicted resonance list (0.527) (Figure 3A). Compared to the binary polarization transfer model, all three spectra experience increased BPHON scores of at least 0.12 (Figure 2A). This is visually illustrated between two simulated spectra with (Figure 3C) and without (Figure 3B) the improved spin physics model used to calculate peak intensity. The simulated spectrum calculated with the improved transfer model described in Eq. 1 has a BPHON score of 0.737 (Figure 3C), while the spectrum calculated using the binary spin physics model has a score of 0.563 (Figure 3B), offering a BPHON score improvement of 0.174.

**Figure 3.**
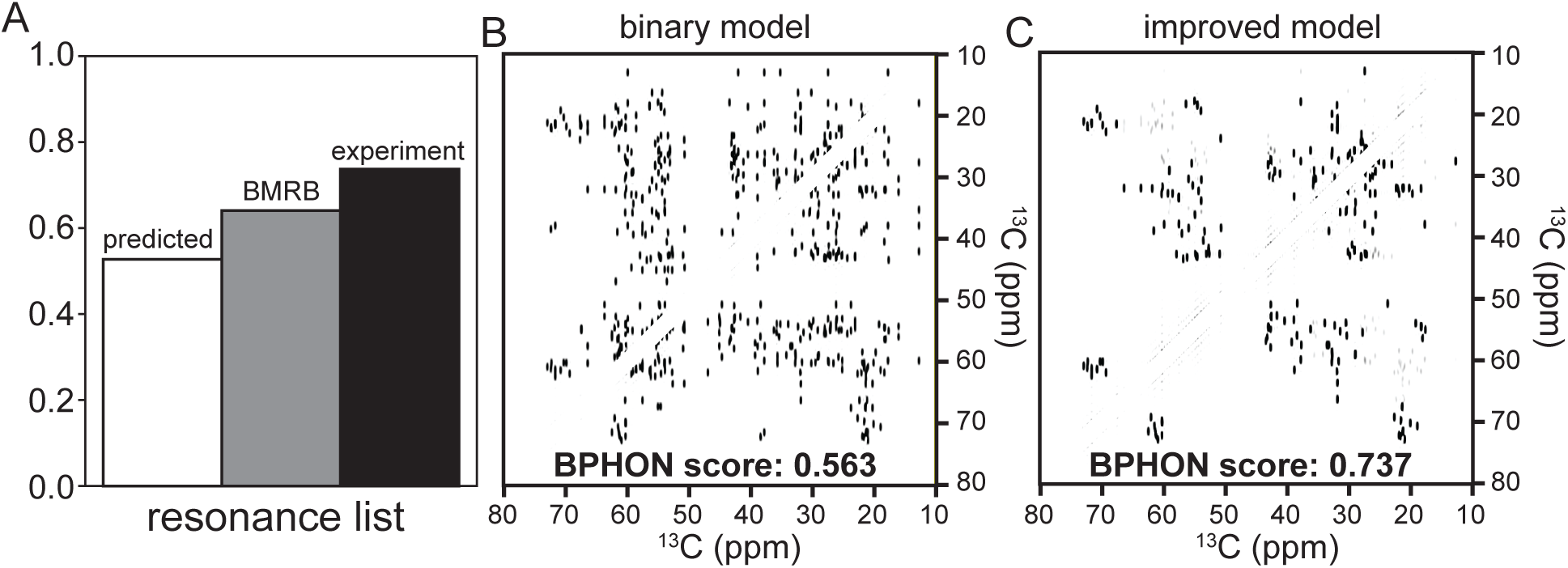
Calculating peak intensity for spectral simulations. A) Histogram of BPHON scores when simulated spectra were calculated with SHIFTX2 (white), BMRB (17810, gray), and experimental (black) resonance lists. B and C) Aliphatic simulated spectra of ^13^C-^13^C spectra with a binary polarization transfer model (B) and with an improved polarization transfer model from equation 1 (C). When a polarization transfer model is used to simulate spectra, the BPHON score improves by 0.171. Simulated spectra did not employ T_2_ relaxation rates and scalar couplings to implement lineshapes. Simulated spectra were scored against an experimental ^13^C-^13^C (DARR, 50 ms) spectrum collected at 14.1 T spinning at 26.6 kHz. Simulated and experimental spectra were processed with a sine bell offset of 59° in both dimensions.

While peak intensities are proportional to the distance between ^13^C atoms, lineshapes provide fruitful information on structural heterogeneity and are critical to recapitulate when simulating protein spectra. This can be described by T_2_ relaxation, which is an inherent property of each sample, depending on the molecular dynamics timescales, as well as the magnetic field, MAS rate and decoupling conditions, contributing together to the homogeneous linewidth (measured in Hz) equivalent to 1/(π*T_2_). Additionally, samples may exhibit inhomogeneous broadening indicated by a characteristic T_2_* value. The spectra simulated for Fig. 2 and 3 were calculated assuming a small homogeneous linewidth (1 Hz), corresponding to a T_2_ > 300 ms. This represents an unrealistically narrow linewidth for a strongly coupled microcrystalline protein, so next we revised these assumptions with site-specific T_2_ relaxation times typically observed for methyl (LW = 20 Hz, T_2_ ∼16 ms), methylene (LW = 50 Hz, T_2_ ∼6 ms), methine (LW = 50 Hz, T_2_ ∼6 ms), and carbonyl (LW = 30 Hz, T_2_ ∼11 ms) groups in proteins (Figure 3A). BPHON also employs known one-bond J-couplings of one-bond sp3-sp3 (35 Hz), sp2-sp2 (75 Hz), and ^13^Cα-^13^C’ (55 Hz) ^13^C-^13^C bonds (Fig 3A).^30^ Linewidths and scalar couplings of ^13^C atoms for the amino acids employed by BPHON are listed in Tables S3 and S4. Three ^13^C-^13^C spectra were then simulated with SHIFTX2, BMRB (17810), and experimental resonance lists and scored against an experimental ^13^C-^13^C (DARR, 50 ms) spectrum. Consistent with the score trend in Figures 2A and 3A, the simulated spectrum calculated with the experimental resonance list has the highest score (0.848), followed by the BMRB (0.752) and the predicted resonance list (0.633) (Figure 4B).

**Figure 4.**
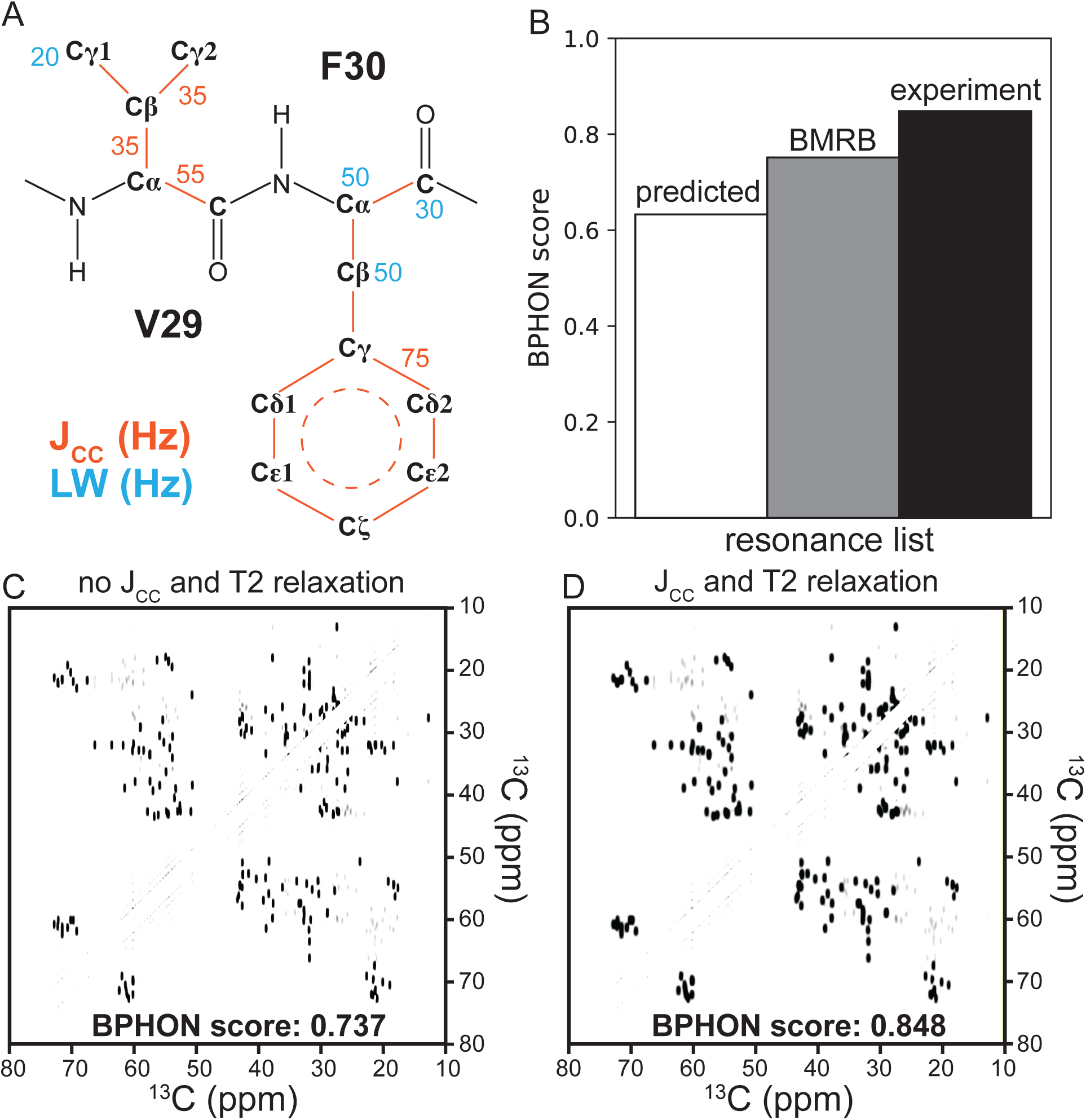
Implementing lineshapes in spectral simulations. A) An example of V29-F30 in microcrystalline GB1 annotated with examples of known one bond ^13^C-^13^C scalar couplings (orange) and linewidths used for C, CH, CH_2_, and CH_3_ atoms. B) Histogram of BPHON scores when simulated spectra were calculated with SHIFTX2 (white), BMRB (17810, gray), and REFINED (black) resonance lists and with T_2_ and J_CC_ values to provide lineshapes. C and D) Simulated spectra of aliphatic region of ^13^C-^13^C spectra with (D) and without (C) T_2_ relaxation rates and J_CC_ values. The BPHON score increases by 0.155 when linewidths and scalar couplings are implemented. Simulated spectra were scored against an experimental ^13^C-^13^C (DARR, 50 ms) spectrum collected at 14.1 T spinning at 26.6 kHz. Simulated and experimental spectra were processed with a sine bell offset of 59° in both dimensions.

However, all three BPHON scores improve at least 0.11 relative to their scores with 1 Hz linewidths (Fig 3A). As a result, the lineshapes in the spectra calculated with T_2_ values more closely approximating microcrystalline GB1 result in an improvement in the BPHON score by 0.111 to a final result of 0.848 (Figure 4C).

The next benchmark assessment depends upon processing. When processing NMR data, the choice of apodization is judiciously chosen as a compromise between sensitivity and resolution. Here, we use a sine bell apodization function to process NMR spectra with 16 different coefficients for the sine bell function, to examine the full range of emphasis between sensitivity and resolution. The adjustment of this coefficient alters the effective line broadening while minimizing the truncation artifacts that would be observed with exponential broadening alone. BPHON scores of spectra simulated with the predicted (squares), BMRB (triangles), and experimental (circles) resonance lists are shown in Figure 5A. As expected, the spectra simulated with the experimental resonance list consistently provide higher BPHON scores than spectra simulated with the BMRB and predicted resonance lists regardless of how the data are processed. However, each spectral series exhibits different trajectories of BPHON scores as a function of effective apodized linewidth. The spectral series simulated with the experimental resonance list has a BPHON score of ∼0.883 from linewidths of ∼80-67 Hz and then steadily drops to 0.731 at a linewidth of 55 Hz, while the BMRB spectral series has a BPHON score of 0.833 from linewidths of 80-73 Hz and then steadily drops to 0.588 at a linewidth of 55 Hz. The predicted spectral series has a BPHON score of 0.762 and rapidly drops to 0.435 at a linewidth of 55 Hz. BPHON scores of spectra simulated with the experimental resonance list exhibits the least change in BPHON scores as a function of how the data are processed. Overlayed simulated (red, simulated with experimental chemical shifts) and experimental (black) Ala CA-CB cross-peaks with average linewidths of 80 Hz (Figure 5B) and 55 Hz (Figure 5C) illustrate this effect. Simulated spectra calculated with less accurate assignments will therefore experience less peak overlap with the experimental spectrum when processed to achieve narrower linewidth.

**Figure 5.**
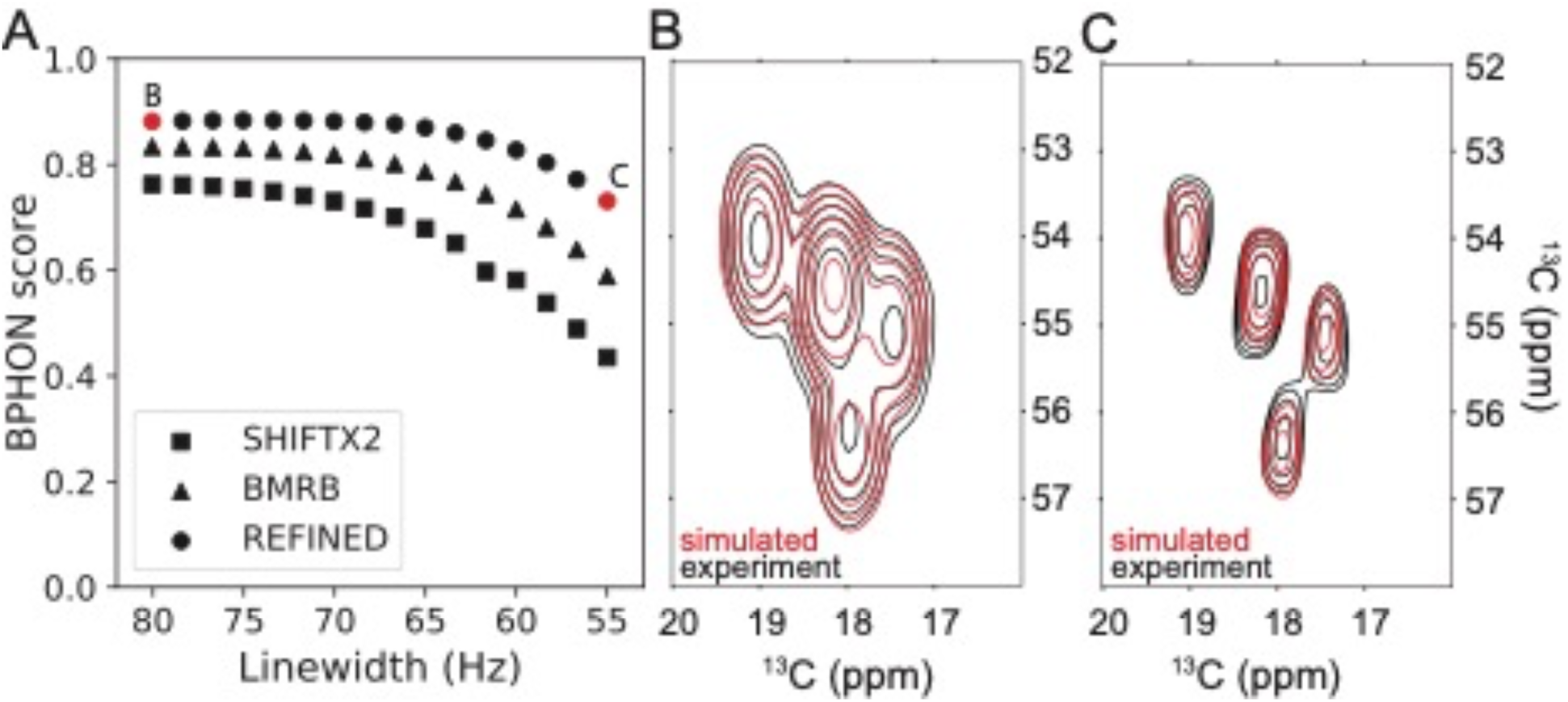
Impact of data processing on BPHON scores. A) scatter plot of BPHON scores vs linewidth (Hz) of spectra processed with 16 different sine bell apodization offsets using SHIFTX2 (squares), BMRB (triangles), and REFINED (circles) resonance lists. For each score, simulated and experimental spectra were processed with matched apodization functions. B and C) Contour overlay between experimental (black) and simulated (red) alpha helical Ala CA-CB cross-peaks with 190 Hz (B) and 147 Hz (C) linewidths. The average linewidth of the four alpha helical Ala peaks were measured and used as the x-axis in part A.

Next, we consider an approach to resonance assignment validation and assess its impact on the results. BPHON offers the capability to generate a hybrid resonance list using both experimental and computational approaches. For example, if an incomplete experimental resonance list is available, BPHON supports using predicted ^13^C chemical shifts when experimental shifts are not available. In the following example 0, 10, 20, 30, 40, 50, 60, 70, 80, 90, and 100% of the ^13^C chemical shifts in the experimental resonance list were randomly removed and replaced with ^13^C shifts predicted by SHIFTX2 (Figure 6). When no experimental ^13^C shifts were used to simulate spectra, the BPHON score was 0.633. As more experimentally derived ^13^C chemical shifts are used to simulate spectra, the BPHON scores steadily increase to 0.848. Therefore, it is recommended that experimental NMR data be used whenever possible and predicted chemical shifts be used to fill in missing experimental chemical shifts. We envision expanding BPHON’s capabilities to support other chemical shift predictors as well as computational programs that leverage quantum mechanics to calculate chemical shifts to fill in missing chemical shifts when experimental shifts are not available.

**Figure 6.**
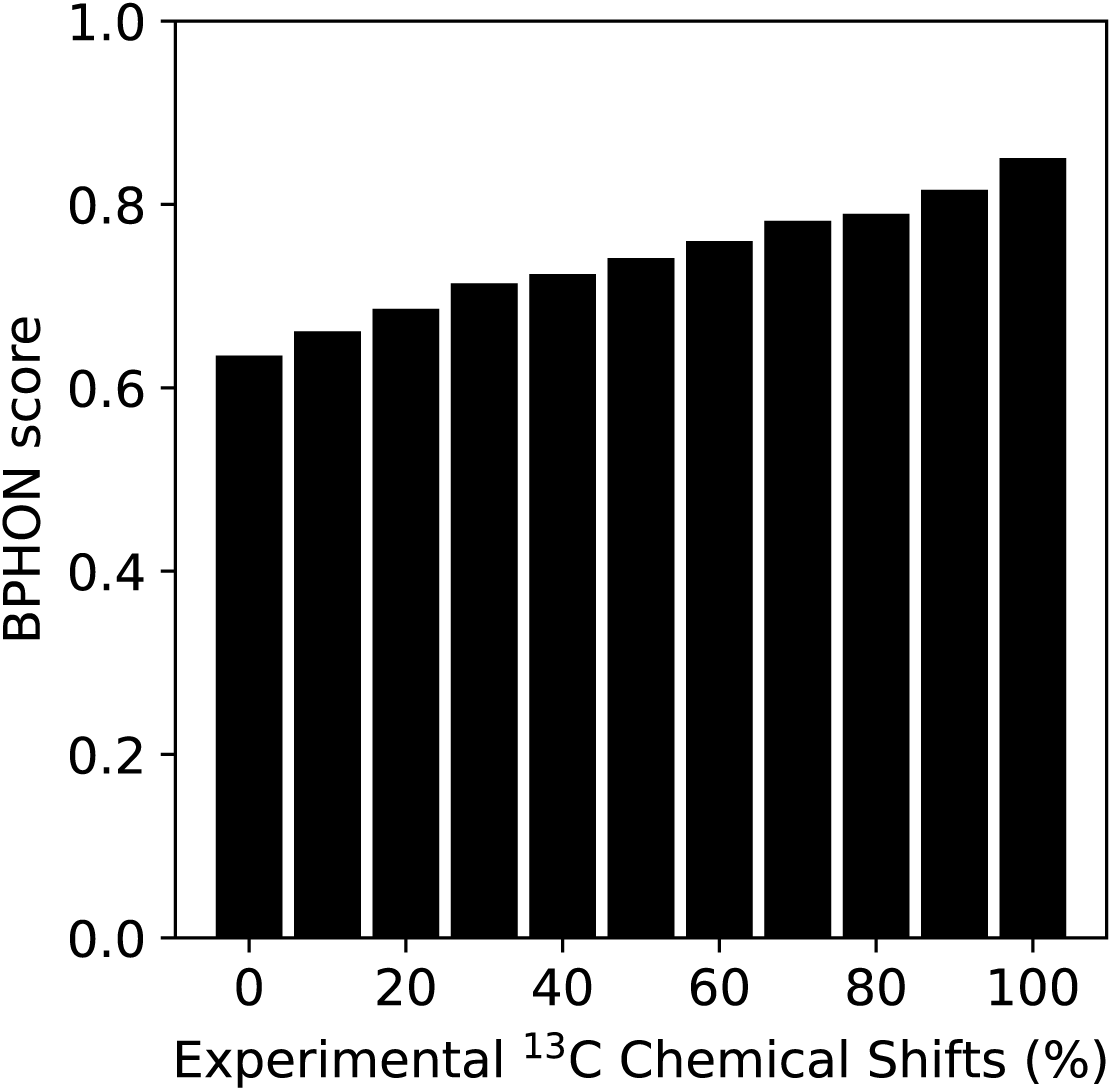
Hybrid resonance lists. Histogram of spectra simulated with hybrid resonance lists with 0-100% experimental ^13^C chemical shifts in 10% increments. Simulated and experimental spectra were processed with 59° sine bell apodization functions.

Finally, we consider two protein candidate structures and use only predicted chemical shifts to simulate and score spectra using BPHON. Model 1 (Figure 7A) is microcrystalline GB1 (2LGI) and model 2 (Figure 7B) is microcrystalline GB1 calculated with only a subset of the restraints deposited to the PDB. The BPHON scores for models 1 and 2 are 0.641 and 0.548, respectively (Figure 7C). Interestingly, the ground truth model (Figure 7A) yields a higher BPHON score than model 2 (Figure 7C) despite using only predicted chemical shifts. While this is not a substitute for assignments, this illustrates that BPHON can provide initial insight into structural accuracy of multiple protein candidate models without any experimental chemical shifts and may serve as an opportunity to guide NMR peak assignments.

**Figure 7.**
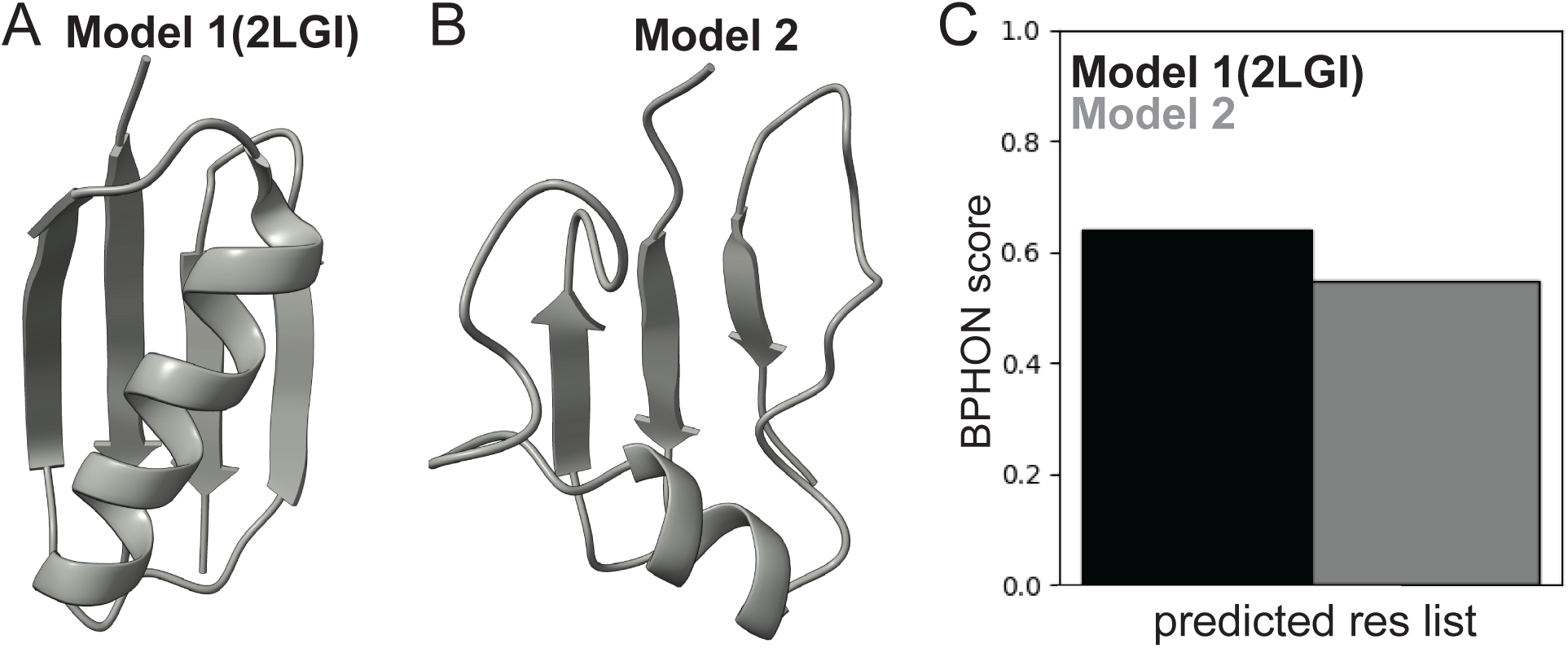
Scoring protein structure accuracy. A and B) model 1 (2LGI) and model 2 used to simulate and score spectra. Model 2 is the lowest energy structure from a GB1 structure calculation with a subset of the restraints submitted to the PDB. C) Histogram of BPHON scores of GB1 models 1 (black) and 2 (gray) simulated with a predicted (SHIFTX2) resonance list. Simulated spectra were scored against an experimental ^13^C-^13^C (DARR, 250 ms) spectrum collected at 14.1 T spinning at 26.6 kHz. Simulated and experimental spectra were processed with a sine bell offset of 59° in both dimensions.

### Graphical User Interface

BPHON is packaged as a plugin to the widely used molecular visualization tool UCSF ChimeraX. BPHON is accessed by clicking on the “Structure Analysis” option under the “Tools” button in the ChimeraX toolbar and the graphical user interface (GUI) panel will then appear on the panel next to the protein structure(s) (Figure 6). Prior to running BPHON, one or multiple protein structures must be open and selected in ChimeraX. The first prompt to be filled by the user is the “Output path” (Figure 8A) and is the directory where the “Experiment name” (Figure 8B) folder is saved, which contains all data associated with the simulation, including the PDB structure, resonance list, NMRPipe simulation and processing scripts, the FID, and the processed simulated spectrum. Next, the user must choose an “Experimental spectrum” (Figure 8C) which is the NMR spectrum that the simulated spectrum will be scored against. The user will then input a resonance list to simulate the spectra. BPHON offers the unique capability to create a hybrid resonance list from a primary, secondary, and tertiary resonance list (Figure 8D). This allows highly confident, but incomplete resonances in the primary resonance list to be preserved while providing missing resonances with in the secondary and tertiary resonance lists. For example, an incomplete experimental resonance list can be used as the primary resonance list and missing resonances can be predicted. Currently, BPHON supports resonance lists in NMR-STAR, NMR exchange format (NEF), and NMRFAM-SPARKY resonance lists. Additionally, BPHON currently supports the chemical shift prediction algorithm SHIFTX2 to predict chemical shifts if none are available. The hybrid resonance list will be saved in the “Experiment name” directory. The user will then decide to simulate ^13^C-^13^C, ^15^N-^13^Cα, or ^15^N-^13^C’ spectra (Figure 8E). For ^13^C-^13^C spectra, the user has the option to remove the principal diagonal as well as spinning side bands to ameliorate effects on the ZNCC score (Fig 8G). If SHIFTX2 is employed to predict chemical shifts, the user can specify the pH and temperature under which to predict the shifts to best match experimental conditions (Figure 8F). Finally, the user defines if/how the simulated and experimental spectra are scored using ZNCC or RMSD. If a scoring metric is selected, a histogram of the BPHON score for each structure will be presented. All spectra can also automatically be displayed in NMRFAM-Sparky.

**Figure 8.**
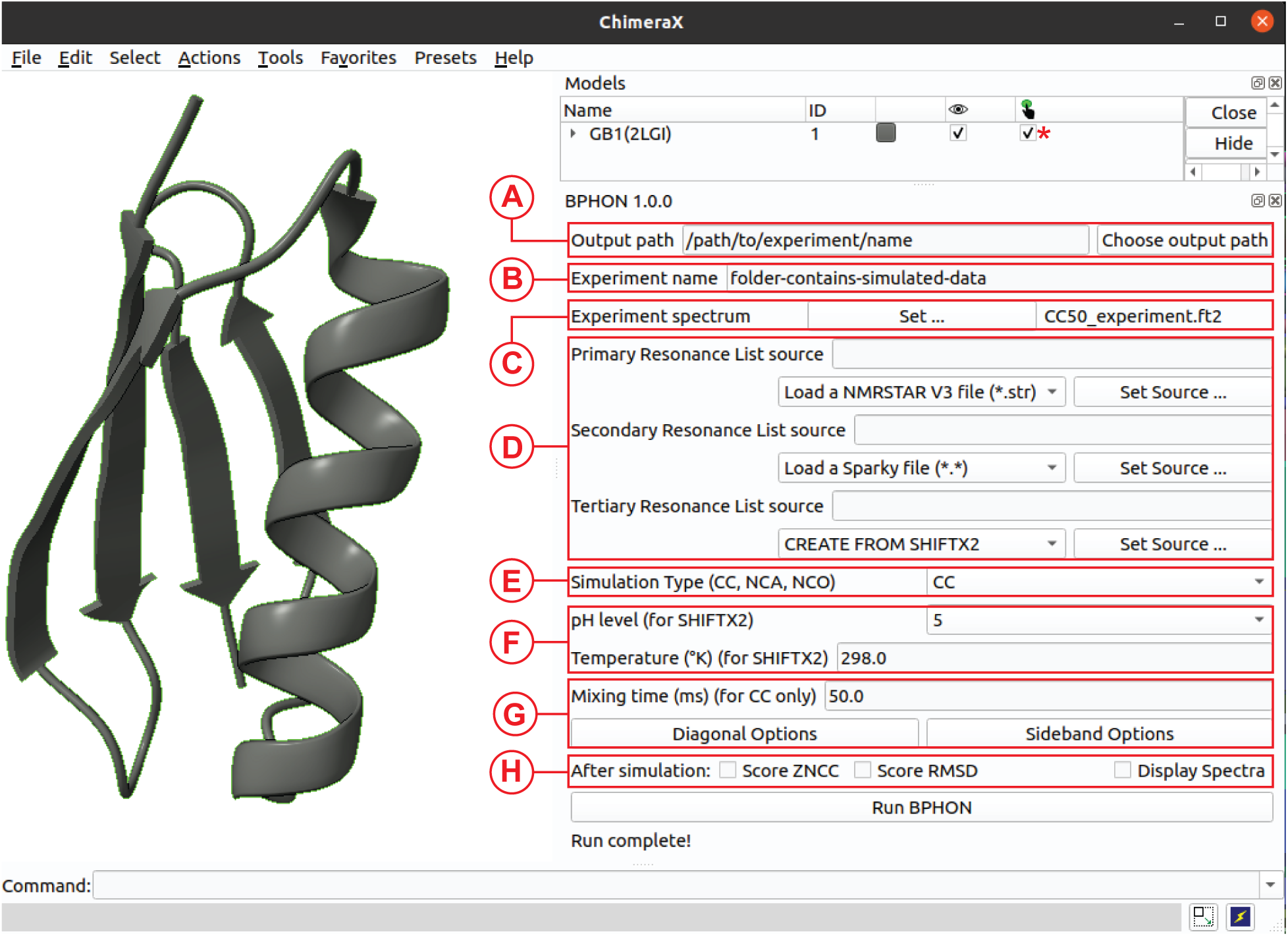
BPHON graphical user interface. The program is a plugin for molecular visualization tool, ChimeraX.

## CONCLUSIONS

BPHON represents a significant advance in NMR-based protein structure validation by integrating spectral simulation with image analysis techniques. Unlike traditional bottom-up approaches that require extensive resonance assignments, our top-down method directly compares experimental spectra with simulated counterparts generated from candidate protein structures. Through rigorous benchmarking, we demonstrated that a spin physics model, incorporating distance-dependent polarization transfer intensity calculations and realistic lineshape modeling, improves scoring accuracy by more than twofold compared to binary models. Assignments are not required but can be used if available. Further improvements to the polarization transfer modeling, for example through explicit numerical simulation with programs such as Spinach,^31^ are anticipated.

The strength of BPHON lies in its flexibility and accessibility. It can operate without experimental chemical shifts while still providing meaningful structural discrimination, though performance improves substantially when experimental data are incorporated. This makes BPHON potentially valuable for challenging systems like large proteins, membrane proteins, and amyloid fibrils, where traditional assignment-based methods often struggle due to spectral complexity and peak overlap. The ability to create hybrid resonance lists combining experimental and predicted chemical shifts offers a pragmatic path forward for partially assigned systems.

We envision BPHON as complementary to existing NMR structure determination workflows, providing early validation of structural models and potentially guiding resonance assignment efforts. Future development will focus on implementing additional spin physics models (like those available in Spinach^31^), alternative image analysis algorithms less sensitive to processing parameters (such as Fourier shell correlation from cryo-EM),^32^ and expanded chemical shift prediction methods including quantum mechanical approaches. By packaging BPHON as a ChimeraX plugin with an intuitive interface, we aim to make sophisticated NMR analysis accessible to the broader structural biology community, potentially accelerating structure determination for challenging protein systems that have previously resisted conventional approaches.

## MATERIALS AND METHODS

### SSNMR Spectroscopy

Adiabatic polarization transfer is described using the shorthand notation as follows:

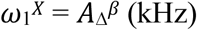

where ω_1_^X^ is the channel specific field strength (e.g. ^1^H, ^15^N, ^13^C), A is the set field strength, β is the shape of the adiabatic ramp, and Δ is the ramp size.

GB1 was prepared according to Franks et al JACS 2005 and packed into a 1.6 mm rotor. Nine ^13^C-^13^C spectra (DARR, t_mix_= 12.5, 25, 50, 100, 150, 200, 250, 375, 500 ms) were collected at 14.1 T (600 MHz ^1^H frequency) equipped with a Balun T3 probe in HCN mode spinning at 26.6 kHz at a sample temperature of −5 +/-5 °C. The ^1^H and ^13^C pulse widths were 2.15 and 2.1 us, respectively. CP transfer from ^1^H to ^13^C occurred such that *ω*_1_^H^ = 97 kHz and *ω*_1_^C^ = 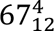 kHz with a contact time of 2.5 ms. SPINAL decoupling at 99 kHz was employed and optimized^33^ with a 5.2 us π pulse during acquisition (15 ms). The spectra were collected with 4 scans and a delay time of 2 s.

### Spectral Simulations

Spectral simulations were performed on a Ryzan workstation with 32 cores running Ubuntu 20.0.4. The spectral simulations were performed using BPHON (version 1.0.0) which is available to download for the program ChimeraX (version 1.6 or later). BPHON requires installation of NMRPipe^34^ (version 10.9 or later) and SHIFTX2 (version 1.10A or later). All BPHON scores presented here are products of scoring both simulated and experimental spectra from 0 to 200 ppm in each ^13^C dimension. Simulated and experimental spectra must be the same size for ZNCC scoring to function. BPHON is also available to use on NMRBox.

### XPLOR-NIH Structure Calculations

Structure calculations were carried out using XPLOR-NIH^35^ version 3.6.6. An extended structure of GB1 was generated from the primary sequence and used as the starting structure. The simulated annealing calculation was comprised of an initial minimization in torsion angle space, a high-temperature step with the bath temperature set to 3,500 K wherein torsion angle dynamics were allowed to proceed for 20 ps or 10,000 steps, whichever came first. Finally, a cooling step was performed where the temperature was ramped down from 3,500 K to 20 K in 50 K increments. At each temperature step during the cooling phase, torsion angle dynamics were carried out for 0.1 ps or 100 steps, whichever came first. Following the cooling phase, a final torsion angle and cartesian minimization were performed for 500 steps each before writing out the PDB files. In addition to canonical bond distance, bond angle and improper angle potentials, a VDW-like repulsive potential was used with a scaling coefficient ramped from 0.004 to 4 over the course of the cooling step. Database potentials were applied to all simulations including a torsion angle database potential with a scaling coefficient ramped from 0.002 to 1 during the cooling phase, a gyrational volume potential scaled from 0.001 to 1.25, and a residue affinity potential scaled from 0.01 to 1.25.

## Supporting information

Software manual

## ASSOCIATED CONTENT

### Data Availability

BPHON is available via git at https://git.doit.wisc.edu/nmrfam-public/bphon-releases.

### Supporting Information

Supporting Information is available free of charge.

## AUTHOR INFORMATION

### Author Contributions

BDH and CMR designed the project, collected and analyzed the data, and wrote the manuscript. BDH, BDZ, RG, and ZH wrote the code for BPHON. FD supported development and provided numerous insightful recommendations and source code from NMRPipe. All authors reviewed and revised the manuscript.

### NOTES

Authors declare no competing financial interest. The authors have files and invention disclosure with the Wisconsin Alumni Research Foundation.

## ACKNOWLEDGMENTS

Authors would like to thank Professor Hannah Wayment-Steele for productive discussion and conceptualization, Dr. Songlin Wang for assistance in the data collection, and Dr. Collin Borcik, Owen Warmuth, and Moses Milchberg for helpful discussions.

## ABBREVIATIONS

CP: cross polarization;
MAS: magic angle spinning

---

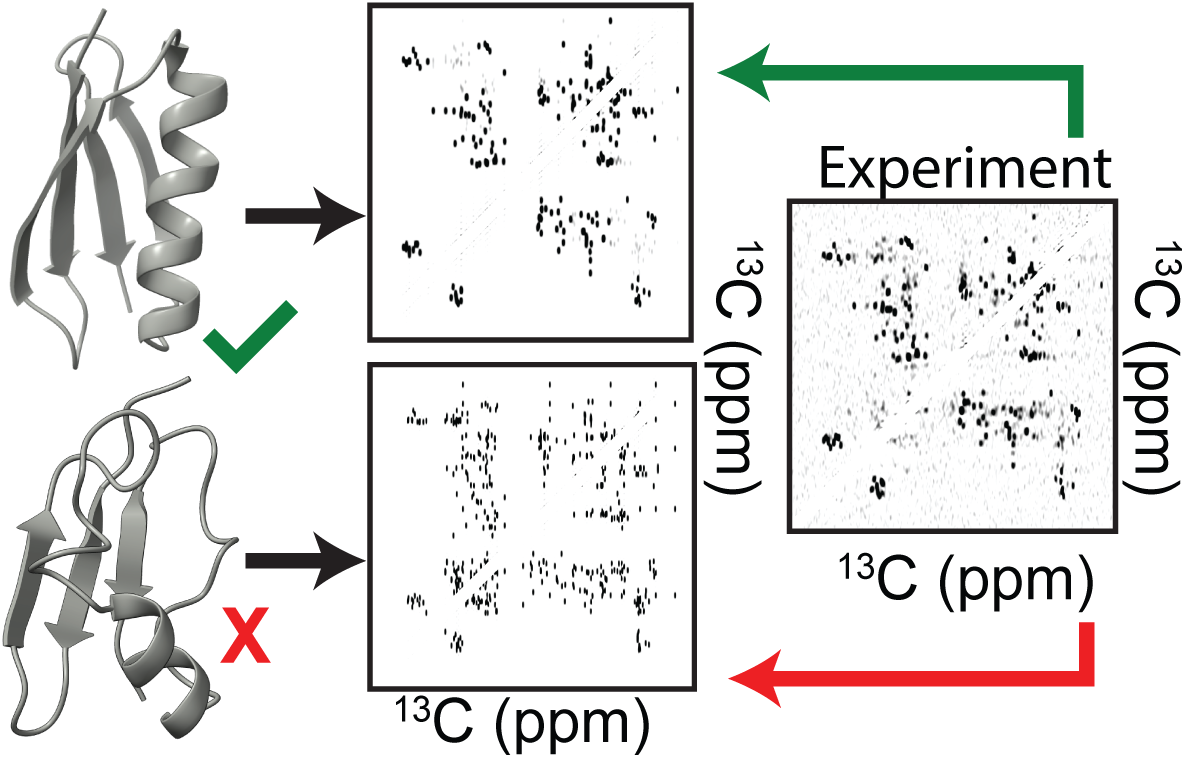

