## Supporting Information for "Top-Down Scoring of Spectral Fitness by Image Analysis for Protein Structure Validation"

Benjamin D. Harding^1,3^, Barry DeZonia^2^, Rajat Garg^1^, Ziling Hu^1^, Katherine Henzler-Wildman^1,2^, Frank Delaglio^5^, Tim Grant^1,*^, Chad M. Rienstra^1,2,4,^*

^1^Department of Biochemistry, University of Wisconsin-Madison, Madison, WI, 53706 USA

^2^National Magnetic Resonance Facility at Madison, University of Wisconsin-Madison, Madison, WI, 53706 USA

^3^Biophysics Graduate Program, University of Wisconsin-Madison, Madison, WI, 53706 USA

^4^Integrated Program in Biochemistry, University of Wisconsin-Madison, Madison, WI, 53706 USA

^5^Morgridge Institute for Discovery, University of Wisconsin-Madison, Madison, WI, 53706 USA

^6^Institute for Bioscience and Biotechnology Research, National Institute of Standards and Technology and the University of Maryland, Rockville, MD, 20850 USA

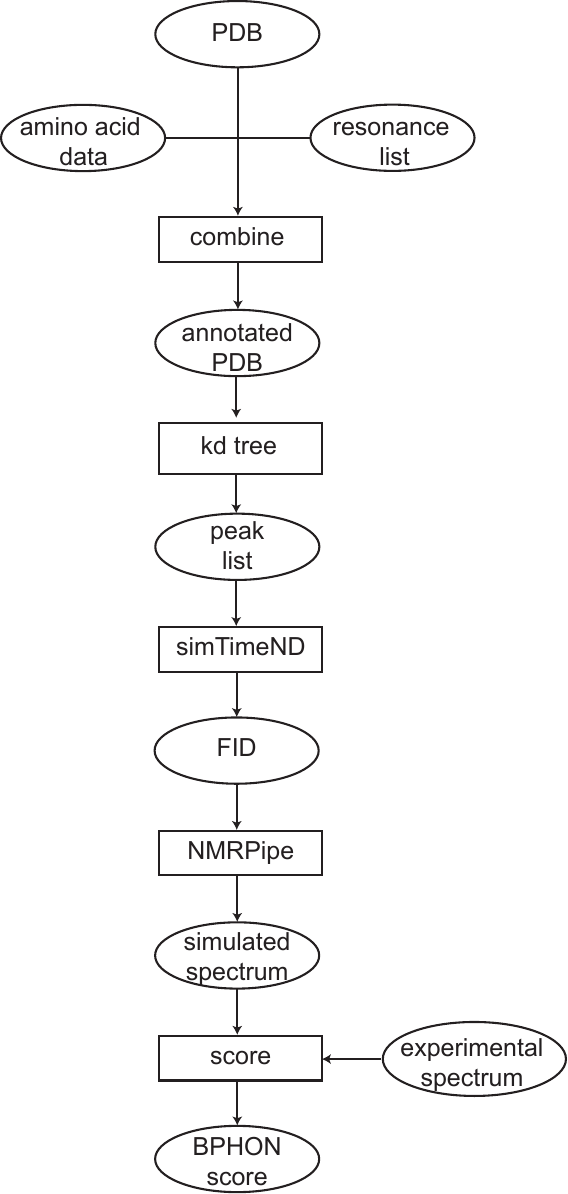
**Figure S1. BPHON pipeline flowchart**. Ovals are inputs/outputs and rectangles are actions. The initial inputs by the user are a protein structure and a resonance list. Additionally, amino acid data, or hard-coded libraries of linewidths and scalar couplings are combined to create an annotated PDB. The PDB is then fed into a k-dimensional tree (kd tree) to generate a peak list which contains the atoms used to construct the peak, linewidths, scalar couplings, and peak height. The peak list is then the input to NMRPipe’s program simTimeND, which appends header information from an experimental NMRPipe conversion script to the peak list to calculate a free induction decay (FID). The FID is then automatically processed using a default NMR processing script to generate an NMR spectrum in the frequency domain. The simulated and experimental spectra are then used as inputs to score their similarity using image analysis, yielding a final BPHON score.

| **experiment type** | **mixing time (ms)** | **cutoff distance (Å)** |
| --- | --- | --- |
| ^13^C-^13^C | 0-50 | ≤4 |
| ^13^C-^13^C | 51-100 | ≤6 |
| ^13^C-^13^C | 101-150 | ≤8 |
| ^13^C-^13^C | 151-200 | ≤10 |
| ^13^C-^13^C | >200 | ≤12 |
| ^15^N-^13^Cα | - | 1.6 |
| ^15^N-^13^CO | - | 1.6 |

**Table S1**. Default guidelines employed by BPHON for defining cutoff distances when constructing ^13^C-^13^C, ^15^N-^13^Cα, and ^15^N-^13^CO peak lists of peptides.

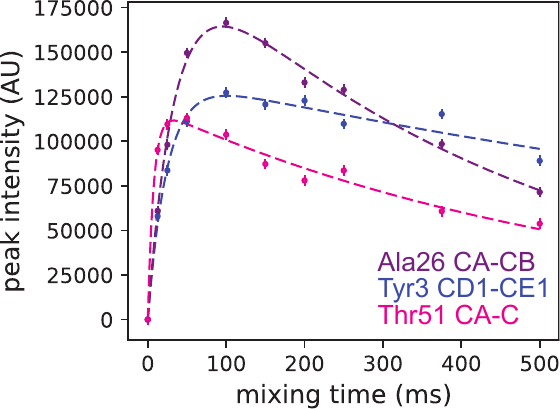

**Figure S2**. Examples of three spins systems with different buildup rates used to fit to equation 1 in the main text. Ala26CA-CB, Tyr3 CD1-CE1, and Thr CA-C are spin systems that are representative of aliphatic, aromatic, and carbonyl spin systems.

| **Peak Position (w1)** | **Peak Position (w2)** | **I_ij_** | **a** | **b** |
| --- | --- | --- | --- | --- |
| Aliphatic (0-80 ppm) | Aliphatic  (0-80 ppm) | 1 | 250 | 0.5 |
| Aliphatic  (0-80 ppm) | Aromatic  (100-165 ppm) | 1 | 50 | 0.5 |
| Cα | Carbonyl  (160-200 ppm) | 1 | 3500 | 0.5 |
| Aliphatic  (0-80 ppm) | Carbonyl  (160-200 ppm) | 1 | 5000 | 0.5 |
| Aromatic  (100-165 ppm) | Aromatic  (100-165 ppm) | 0.5 | 250 | 0.5 |
| Aromatic  (100-165 ppm) | Carbonyl  (160-200 ppm) | 1 | 50 | 0.5 |
| Carbonyl  (160-200 ppm) | Carbonyl  (160-200 ppm) | 1 | 5000 | 0.5 |

**Table S2**. Table of parameters from equation 1 in the main text used calculate peak intensity based on location of the chemical shift.

| amino acid | ^13^C atom | linewidth (Hz) |
| --- | --- | --- |
| ALA | CA | 50 |
| ALA | CB | 20 |
| ALA | C | 30 |
| ARG | CA | 50 |
| ARG | CB | 50 |
| ARG | CG | 50 |
| ARG | CD | 50 |
| ARG | CZ | 30 |
| ARG | C | 30 |
| ASN | CA | 50 |
| ASN | CB | 50 |
| ASN | CG | 30 |
| ASN | C | 30 |
| ASP | CA | 50 |
| ASP | CB | 50 |
| ASP | CG | 30 |
| ASP | C | 30 |
| ASX | CA | 50 |
| ASX | CB | 50 |
| ASX | CG | 30 |
| ASX | C | 30 |
| CYS | CA | 50 |
| CYS | CB | 50 |
| CYS | C | 30 |
| GLN | CA | 50 |
| GLN | CB | 50 |
| GLN | CG | 50 |
| GLN | CD | 30 |
| GLN | C | 30 |
| GLU | CA | 50 |
| GLU | CB | 50 |
| GLU | CG | 50 |
| GLU | CD | 30 |
| GLU | C | 30 |
| GLY | CA | 50 |
| GLY | C | 30 |
| HIS | CA | 50 |
| HIS | CB | 50 |
| HIS | CG | 30 |
| HIS | CD2 | 50 |
| HIS | CE1 | 50 |
| HIS | C | 30 |
| ILE | CA | 50 |
| ILE | CB | 50 |
| ILE | CG1 | 50 |
| ILE | CG2 | 20 |
| ILE | CD1 | 20 |
| ILE | C | 30 |
| LEU | CA | 50 |
| LEU | CB | 50 |
| LEU | CG | 50 |
| LEU | CD1 | 20 |
| LEU | CD2 | 20 |
| LEU | C | 30 |
| LYS | CA | 50 |
| LYS | CB | 50 |
| LYS | CG | 50 |
| LYS | CD | 50 |
| LYS | CE | 50 |
| LYS | C | 30 |
| MET | CA | 50 |
| MET | CB | 50 |
| MET | CG | 50 |
| MET | CE | 20 |
| MET | C | 30 |
| PHE | CA | 50 |
| PHE | CB | 50 |
| PHE | CG | 30 |
| PHE | CD1 | 50 |
| PHE | CD2 | 50 |
| PHE | CE1 | 50 |
| PHE | CE2 | 50 |
| PHE | CZ | 50 |
| PHE | C | 30 |
| PRO | CA | 50 |
| PRO | CB | 50 |
| PRO | CG | 50 |
| PRO | CD | 50 |
| PRO | C | 30 |
| SER | CA | 50 |
| SER | CB | 50 |
| SER | C | 30 |
| THR | CA | 50 |
| THR | CB | 50 |
| THR | CG2 | 20 |
| THR | C | 30 |
| TRP | CA | 50 |
| TRP | CB | 50 |
| TRP | CG | 30 |
| TRP | CD1 | 50 |
| TRP | CD2 | 30 |
| TRP | CE2 | 30 |
| TRP | CE3 | 50 |
| TRP | CN2 | 50 |
| TRP | CH2 | 50 |
| TRP | CZ2 | 50 |
| TRP | CZ3 | 50 |
| TRP | C | 30 |
| TYR | CA | 50 |
| TYR | CB | 50 |
| TYR | CG | 30 |
| TYR | CD1 | 50 |
| TYR | CD2 | 50 |
| TYR | CE1 | 50 |
| TYR | CE2 | 50 |
| TYR | CZ | 30 |
| TYR | C | 30 |
| VAL | CA | 50 |
| VAL | CB | 50 |
| VAL | CG1 | 20 |
| VAL | CG2 | 20 |
| VAL | C | 30 |

**Table S3**. Linewidths (Hz) dictionary employed by BPHON to assign linewidths to ^13^C atoms in peptides. Methyl, methylene, methine, and carbons with no directly bonded ^1^H have linewidths of 20, 50, 50, and 30 Hz, respectively.

| amino acid | peptide bond | scalar coupling (Hz) |
| --- | --- | --- |
| ALA | CA-C | 55 |
| ALA | CA-CB | 35 |
| ARG | CA-C | 55 |
| ARG | CA-CB | 35 |
| ARG | CB-CG | 35 |
| ARG | CG-CD | 35 |
| ASN | CA-C | 55 |
| ASN | CA-CB | 35 |
| ASN | CB-CG | 35 |
| ASP | CA-C | 55 |
| ASP | CA-CB | 35 |
| ASP | CB-CG | 35 |
| CYS | CA-C | 55 |
| CYS | CA-CB | 35 |
| GLN | CA-C | 55 |
| GLN | CA-CB | 35 |
| GLN | CB-CG | 35 |
| GLN | CG-CD | 35 |
| GLU | CA-C | 55 |
| GLU | CA-CB | 35 |
| GLU | CB-CG | 35 |
| GLU | CG-CD | 35 |
| GLY | CA-C | 55 |
| HIS | CA-C | 55 |
| HIS | CA-CB | 35 |
| HIS | CB-CG | 35 |
| HIS | CG-CD2 | 75 |
| ILE | CA-C | 55 |
| ILE | CA-CB | 35 |
| ILE | CB-CG2 | 35 |
| ILE | CB-CG1 | 35 |
| ILE | CG1-CD1 | 35 |
| LEU | CA-C | 55 |
| LEU | CA-CB | 35 |
| LEU | CB-CG | 35 |
| LEU | CG-CD1 | 35 |
| LEU | CG-CD2 | 35 |
| LYS | CA-C | 55 |
| LYS | CA-CB | 35 |
| LYS | CB-CG | 35 |
| LYS | CG-CD | 35 |
| LYS | CD-CE | 35 |
| MET | CA-C | 55 |
| MET | CA-CB | 35 |
| MET | CB-CG | 35 |
| PHE | CA-C | 55 |
| PHE | CA-CB | 35 |
| PHE | CB-CG | 35 |
| PHE | CG-CD1 | 75 |
| PHE | CG-CD2 | 75 |
| PHE | CD1-CE1 | 75 |
| PHE | CD2-CE2 | 75 |
| PHE | CE1-CZ | 75 |
| PHE | CE2-CZ | 75 |
| PRO | CA-C | 55 |
| PRO | CA-CB | 35 |
| PRO | CB-CG | 35 |
| PRO | CG-CD | 35 |
| SER | CA-C | 55 |
| SER | CA-CB | 35 |
| THR | CA-C | 55 |
| THR | CA-CB | 35 |
| THR | CB-CG2 | 35 |
| TRP | CA-C | 55 |
| TRP | CA-CB | 35 |
| TRP | CB-CG | 35 |
| TRP | CG-CD1 | 75 |
| TRP | CG-CD2 | 75 |
| TRP | CD2-CE2 | 75 |
| TRP | CD2-CE3 | 75 |
| TRP | CE2-CZ2 | 75 |
| TRP | CE3-CZ3 | 75 |
| TRP | CZ2-CH2 | 75 |
| TRP | CZ3-CH2 | 75 |
| TYR | CA-C | 55 |
| TYR | CA-CB | 35 |
| TYR | CB-CG | 35 |
| TYR | CG-CD1 | 75 |
| TYR | CG-CD2 | 75 |
| TYR | CD1-CE1 | 75 |
| TYR | CD2-CE2 | 75 |
| TYR | CE1-CZ | 75 |
| TYR | CE2-CZ | 75 |
| VAL | CA-C | 55 |
| VAL | CA-CB | 35 |
| VAL | CB-CG1 | 35 |
| VAL | CB-CG2 | 35 |

**Table S4**. The dictionary of one-bond scalar couplings (*J_CC_*) employed by BPHON when calculating ^13^C-^13^C NMR spectra of peptides. Hybridized sp3-sp3, sp2-sp2, Cα-C’ have scalar couplings of 35, 75, and 55 Hz, respectively.
