## Supplementary material for "Top-Down Scoring of Spectral Fitness by Image Analysis for Protein Structure Validation": Software manual

**NMRFAM-BPHON**

**Operations Manual**

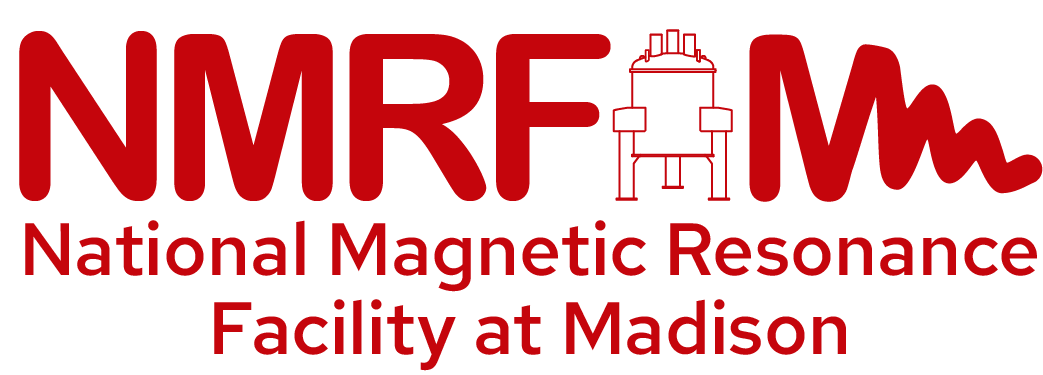

**UW-Madison**

**Version 1.0.0**

### 1. Introduction:

This manual is written for installation of BPHON (1.0.0) and related programs on a Ryzen workstation with 32 cores running Ubuntu 20.04.06.

### 2. Installation:

BPHON requires ChimeraX (1.8 or later), SHIFTX2 (1.13 or later), and NMRPipe (10.9 or later) to be downloaded on your local workstation. Download and install the following programs here:

- ChimeraX (1.8 or later): <https://www.cgl.ucsf.edu/chimerax/download.html>
- NMRPipe (10.9 or later): <https://www.ibbr.umd.edu/nmrpipe/install.html>
- SHIFTX2 (1.13 or later): <http://www.shiftx2.ca/download.html>

Detailed procedures for downloading these programs are available in the Troubleshooting section.

A compressed (tar.gz) BPHON can be downloaded from GitHub at <https://git.doit.wisc.edu/nmrfam/bphon>. Unzip this path in a user-defined directory (this will be called /path/to/bphon/directory/)

a) Open ChimeraX
b) In the ChimeraX command line, execute devel install /path/to/bphon/directory/chimerax-plugin/tut_tool_qt

Successful installation of BPHON will yield the following in the ChimeraX log:

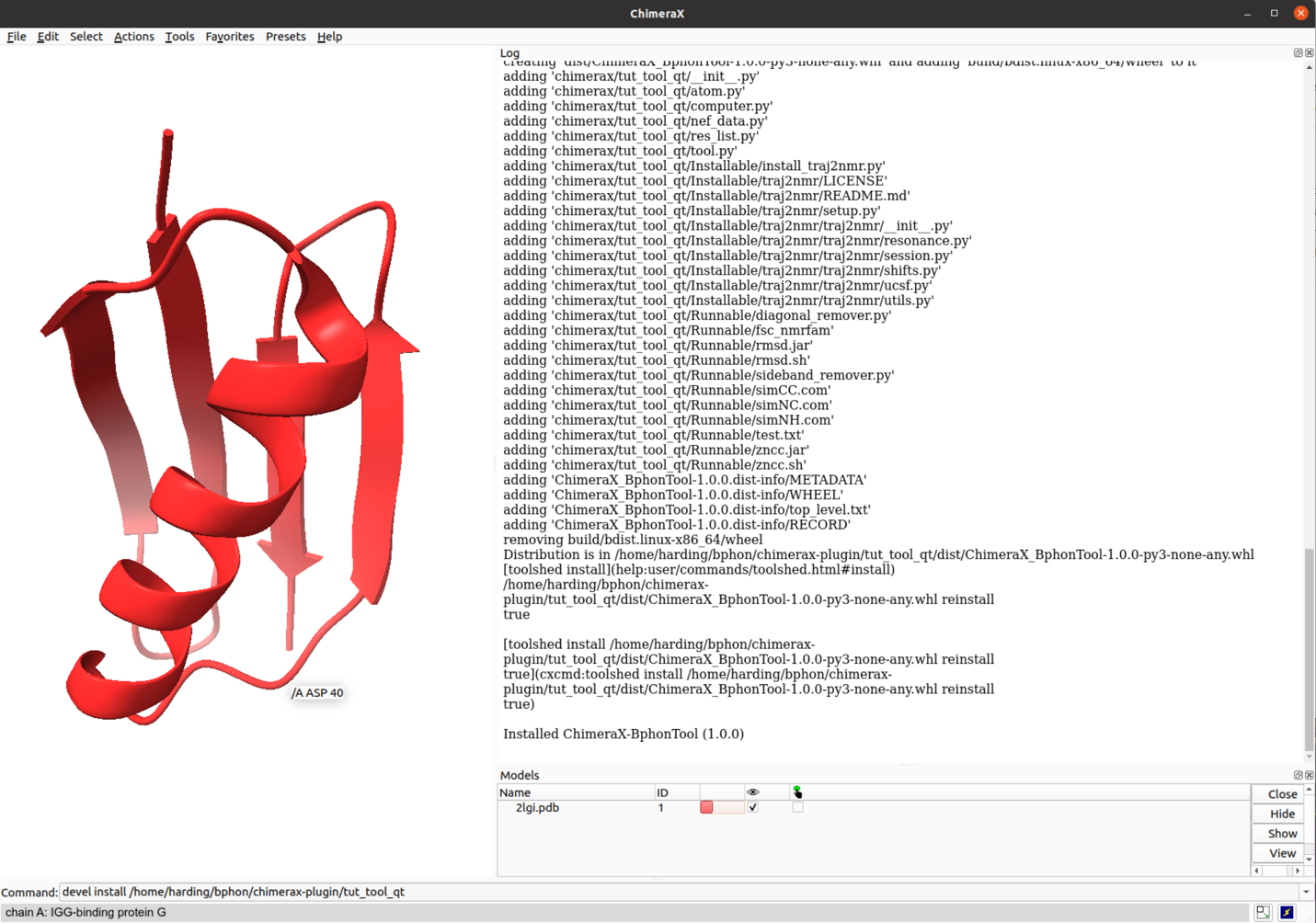

Example command to install bphon:

devel install /home/harding/bphon/chimerax-plugin/tut_tool_qt

ChimeraX Log upon installation will print everything BPHON installs and ends with “**Installed ChimeraX-BphonTool (1.0.0)**”

**Figure 1. ChimeraX example installation command and ChimeraX log upon installation of BPHON (1.0.0)**

### 3. Example Simulation

To simulate a spectrum, open a protein in ChimeraX. For this example, we use the protein GB1 (PDB 2LGI). **Be sure to select your protein**!

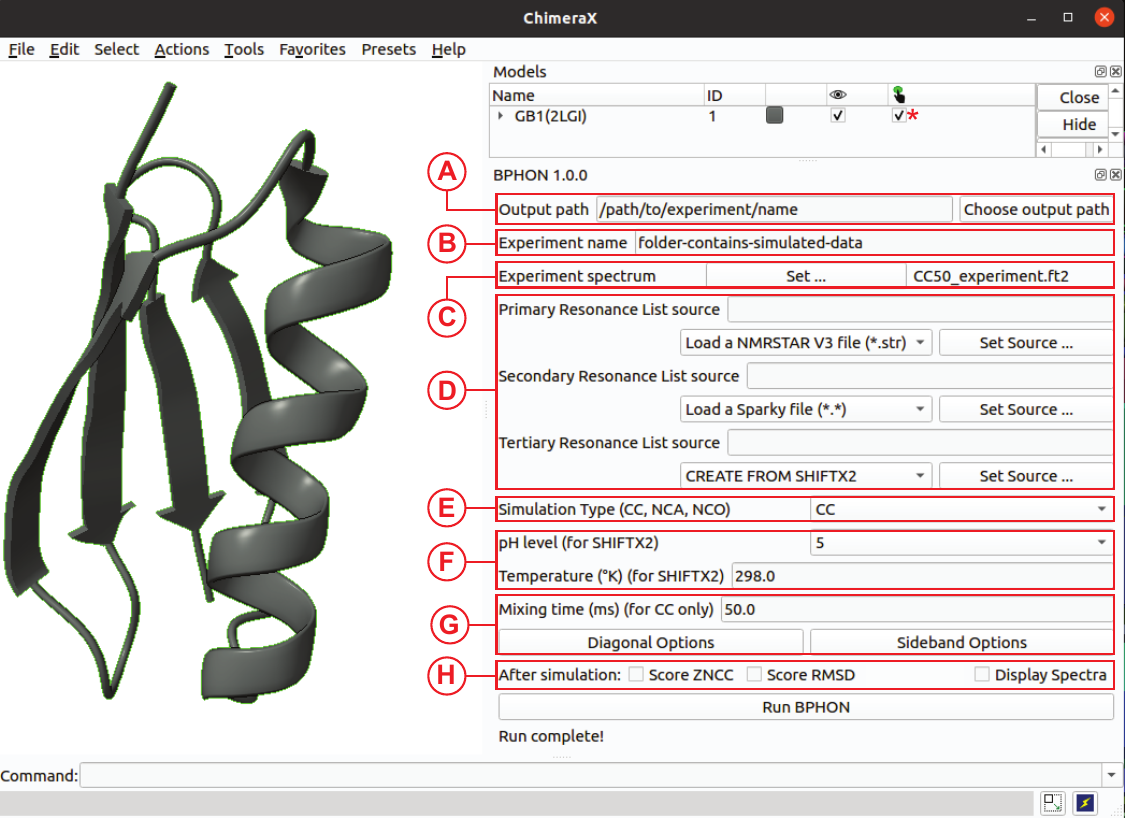

A: Choose a directory path where all the simulated data will be saved.

B: Name of the folder where all simulated data will be stored

C: Choose the experimental file the simulated spectrum will be scored against. We typically use a ^13^C-^13^C (DARR, 50 ms) spectrum collected at 14.1 T spinning at 26.6 kHz processed with a 59˚ sine bell offset (location: /mnt/nmrfam_data/taurus/2022/NOV22/120_CC_DARR_50ms.fid/CC50_experiment.ft2)

D: Resonance list. For this example, just used SHIFTX2 as the primary resonance list

E: Choose the default “CC” as the simulation type (Default)

F: Adjust pH and Temperature. Default values are 5 and 298 K, respectively.

G: Choose the mixing time (50.0 ms Default). Remove principal diagonal and spinning sidebands by typing in magnetic field strength (^1^H Larmour) and spinning rate.

H: Choose “ZNCC” to score simulated and experimental spectra. Choose “Display Spectra” to view overlay of simulated and experimental spectra.

Click “Run BPHON”

### 4. Troubleshooting

This section includes detailed procedures for downloading programs that interact with BPHON, including ChimeraX, NMRPipe, and SHIFTX2.

4.1 Downloading ChimeraX:

a) Go to the **ChimeraX download page**:
<https://www.cgl.ucsf.edu/chimerax/download.html>. Find the **Linux version** and look for **ChimeraX 1.8** (or later).
Alternatively, download it directly using wget:

wget <https://www.cgl.ucsf.edu/chimerax/download/1.8/ChimeraX-1.8-linux_x86_64.tar.gz>

b) Unzip the compressed file: tar -xvzf ChimeraX-1.8-linux_x86_64.tar.gz

4.2 Downloading NMRPipe:

a) we recommend first making a directory for NMRPipe (i.e. /home/Programs/NMRPipe)

b) in terminal: cd /home/Programs/NMRPipe

c) Download the required files from NMRPipe

wget https://www.ibbr.umd.edu/nmrpipe/install.com

wget https://www.ibbr.umd.edu/nmrpipe/binval.com

wget https://www.ibbr.umd.edu/nmrpipe/NMRPipeX.tZ

wget https://www.ibbr.umd.edu/nmrpipe/s.tZ

wget https://www.ibbr.umd.edu/nmrpipe/dyn.tZ

wget https://spin.niddk.nih.gov/bax/software/talos_nmrPipe.tZ

wget https://spin.niddk.nih.gov/bax/software/smile/plugin.smile.tZ

c) Download the following libraries in terminal:

sudo apt-get install tcsh

sudo apt-get install xterm

sudo apt-get install lib32z1

sudo apt-get install libx11-6:i386

sudo apt-get install libxext6:i386

sudo apt-get install xfonts-75dpi

sudo apt-get install msttcorefonts

d) Change shell to tcsh
e) In the NMRPipe directory, execute the ./install script
f) Copy the following initialization scripts into ~/.cshrc

if (-e /usr/local/NMRPipe/com/nmrInit.linux212_64.com) then

source /usr/local/NMRPipe/com/nmrInit.linux212_64.com

endif

if (-e /usr/local/NMRPipe/dynamo/com/dynInit.com) then

source /usr/local/NMRPipe/dynamo/com/dynInit.com

endif

if (-e /usr/local/NMRPipe/com/font.com) then

source /usr/local/NMRPipe/com/font.com

endif

g) In a new terminal window, execute “nmrPipe” and it should return the version of NMRPipe you installed. Example

** NMRPipe System Version 10.9 Rev 2021.258.11.26 64-bit **

4.3 Downloading SHIFTX2:

a) In terminal, execute sudo apt-get install python2.7 default-jre
b) In a user-defined directory, execute wget <http://www.shiftx2.ca/download/shiftx2-v113-linux-20180808.tgz>
c) Execute tar -xzvf shiftx2-v113-linux-20180808.tgz
d) cd shiftx2-linux
e) In the code, change occurrences of “python2.7” to “python” by executing awk '{ gsub(/python/, "python2.7"); print }' shiftx2.py > temp && mv temp shiftx2.py
f) Allow the command to be executable by chmod +x shiftx2.py
g) Add the directory to your PATH variable by executing
   a. export SHIFTX2_DIR=~/Programs/shiftx2-linux
   b. export PATH=$PATH:$SHIFTX2_DIR

4.3 Downloading NMRFAM-SPARKY:

a) Instruction on how to download and install NMRFAM-SPARKY can be found at the following address: <https://nmrfam.wisc.edu/nmrfam-sparky-distribution/>
b) Upon installation, execution of the command “sparky” in Terminal should start the program.
